# A *w*AlbB *Wolbachia* transinfection displays stable phenotypic effects across divergent *Aedes aegypti* mosquito backgrounds

**DOI:** 10.1101/2021.06.25.450002

**Authors:** Perran A. Ross, Xinyue Gu, Katie L. Robinson, Qiong Yang, Ellen Cottingham, Yifan Zhang, Heng Lin Yeap, Xuefen Xu, Nancy M. Endersby-Harshman, Ary A. Hoffmann

**Affiliations:** Pest and Environmental Adaptation Research Group, Bio21 Institute and the School of BioSciences, the University of Melbourne, Parkville, VIC, Australia; CSIRO Land and Water, Black Mountain, Canberra, ACT, Australia

**Keywords:** *Wolbachia*, *w*AlbB, *Aedes aegypti*, cytonuclear interactions, genomics, population replacement, population suppression

## Abstract

*Aedes* mosquitoes harboring intracellular *Wolbachia* bacteria are being released in arbovirus and mosquito control programs. With releases taking place around the world, understanding the contribution of host variation to *Wolbachia* phenotype is crucial. We generated a *Wolbachia* transinfection (*w*AlbB^Q^) in *Aedes aegypti* and performed backcrossing to introduce the infection into Australian or Malaysian nuclear backgrounds. Whole *Wolbachia* genome sequencing shows that the *w*AlbB^Q^ transinfection is near-identical to the reference *w*AlbB genome, suggesting few changes since the infection was first introduced to *Ae. aegypti* over 15 years ago. However, these sequences were distinct from other available *w*AlbB genome sequences, highlighting the potential diversity of *w*AlbB in natural *Ae. albopictus* populations. Phenotypic comparisons demonstrate effects of *w*AlbB infection on egg hatch and nuclear background on fecundity and body size, but no interactions between *w*AlbB infection and nuclear background for any trait. The *w*AlbB infection was stable at high temperatures and showed perfect maternal transmission and cytoplasmic incompatibility regardless of host background. Our results demonstrate the stability of *w*AlbB across host backgrounds and point to its long-term effectiveness for controlling arbovirus transmission and mosquito populations.

## Introduction

*Wolbachia* are intracellular, maternally-inherited bacteria found within about half of all insect species (Weinert *et al*. 2015; Sazama *et al*. 2019). *Wolbachia*-infected *Aedes aegypti* mosquitoes are being released into field populations as a way of controlling mosquito populations and arbovirus transmission (Ross et al. 2019c). Release programs involve strains of *Wolbachia* that have been introduced artificially into *Ae. aegypti*, which do not harbor *Wolbachia* naturally (Gloria-Soria *et al*. 2018; Ross *et al*. 2020b). Most *Wolbachia* infections in mosquitoes induce cytoplasmic incompatibility, where *Wolbachia*-infected males cannot produce viable offspring with females that are not infected (Xi et al. 2005b; McMeniman *et al*. 2009; Walker *et al*. 2011). *Wolbachia* infections can also suppress virus replication within the mosquito, limiting their ability to transmit dengue, Zika and other arboviruses (Terradas and McGraw 2017). *Wolbachia* can spread through target mosquito populations by inducing cytoplasmic incompatibility, which provides a frequency-dependent fitness advantage to *Wolbachia*-infected females (Hoffmann *et al*. 2011). Wolbachia infections can also be used to suppress mosquito populations through cytoplasmic incompatibility (Mains *et al*. 2019; Zheng *et al*. 2019; Crawford *et al*. 2020) as well as deleterious host fitness effects (Ritchie *et al*. 2015).

A variety of *Wolbachia* strains originating from *Drosophila* and mosquitoes have been introduced into *Ae. aegypti* through microinjection (Fraser *et al*. 2017; Ant et al. 2018). These strains show diverse effects on host fitness, virus blocking and cytoplasmic incompatibility (Ross *et al*. 2019c). Population replacement releases currently involve two *Wolbachia* strains: wMel and wAlbB. Both strains have successfully established in natural *Ae. aegypti* populations (Hoffmann *et al*. 2011; Garcia *et al*. 2019; Nazni *et al*. 2019; Tantowijoyo *et al*. 2020) and have similar viral blocking effects (Ant *et al*. 2018; Flores *et al*. 2020). Population replacement programs have demonstrated efficacy against dengue transmission, with fewer dengue cases in areas where *Wolbachia* infections are at a high frequency in the *Ae. aegypti* population (O’Neill *et al*. 2018; Nazni *et al*. 2019; Ryan *et al*. 2019; Indriani *et al*. 2020; Pinto *et al*. 2021). However, establishment success can vary dramatically in different locations, even with the same *Wolbachia* strain. For instance, wMel readily invaded Australian *Ae. aegypti* populations (Hoffmann *et al*. 2011; Schmidt *et al*. 2017; O’Neill *et al*. 2018; Ryan *et al*. 2019) but did not reach high frequencies in many parts of Niterói and Rio de Janeiro in Brazil despite multiple rounds of releases (Gesto et al. 2021; Pinto *et al*. 2021).

*Wolbachia* release success depends on the properties of the *Wolbachia* strain, including host fitness costs, cytoplasmic incompatibility, maternal transmission, virus blocking and environmental stability. The wAlbB *Wolbachia* strain occurs naturally in *Ae. albopictus* and was the first strain transferred to *Ae. aegypti* through microinjection (Xi *et al*. 2005b). *w*AlbB has limited effects on dengue virus transmission in its native host (Lu *et al*. 2012; Mousson *et al*. 2012) but shows strong blocking against a range of arboviruses in *Ae. aegypti* (Bian *et al*. 2010; Joubert *et al*. 2016; Ant *et al*. 2018; Bhattacharya *et al*. 2020) and complete cytoplasmic incompatibility (Xi *et al*. 2005b). *w*AlbB infection induces substantial host fitness costs, including reduced tolerance to starvation (Ross *et al*. 2016; Foo *et al*. 2019) and thermal stress (Ross *et al*. 2019b), decreased quiescent egg viability (Axford *et al*. 2016; Joubert *et al*. 2016) and high rates of female infertility following egg storage (Lau *et al*. 2021). However, *w*AlbB is relatively stable at high temperatures compared to wMel (Ross *et al*. 2017b), prompting its deployment in field trials in Kuala Lumpur, Malaysia. Releases of *w*AlbB led to stable population replacement in some trial locations with a corresponding reduction in dengue transmission (Nazni *et al*. 2019). Aedes *aegypti* carrying *w*AlbB have also been released in population suppression programs that rely on cytoplasmic incompatibility to reduce wild female fertility (Mains *et al*. 2019; Crawford *et al*. 2020).

When a *Wolbachia* strain is released into a new environment, it is important to understand factors that can affect its establishment. *Wolbachia* releases are taking place in genetically divergent mosquito populations (Gloria-Soria et al. 2016; Schmidt *et al*. 2020) and host genotype may influence release outcomes. Host genotype can influence *Wolbachia’s* effects on host fitness (Reynolds *et al*. 2003; Dean 2006; Kyritsis *et al*. 2019; Carvalho *et al*. 2020), cytoplasmic incompatibility (Reynolds and Hoffmann 2002; Bordenstein *et al*. 2003) and viral blocking (Terradas *et al*. 2017; Ford *et al*. 2019). *Wolbachia* releases in new locations often involve backcrossing to introduce the target *Wolbachia* infection into locally-adapted mosquitoes (Dutra *et al*. 2015; Garcia *et al*. 2019; Tantowijoyo *et al*. 2020). Since *Wolbachia* and mitochondria are co-inherited, *Wolbachia* infections carry along their associated mitochondria when they invade (Rasgon *et al*. 2006; Yeap *et al*. 2016), resulting in mosquito populations with foreign mitochondria (Garcia *et al*. 2019; Tantowijoyo *et al*. 2020). Mismatches between mitochondrial and nuclear genotypes in other insects can have severe deleterious effects (Hoekstra *et al*. 2013; Meiklejohn *et al*. 2013; Rank *et al*. 2020). However, such effects have not been tested in mosquitoes.

In this study, we generated a new wAlbB transinfection in *Ae. aegypti* with an Australian mitochondrial haplotype and nuclear background, denoted wAlbB^Q^. We then used antibiotic curing and backcrossing to generate populations to investigate effects of wAlbB infection across two nuclear backgrounds. We found that phenotypic effects were stable across backgrounds, with no evidence for deleterious effects when mitochondrial haplotypes and nuclear backgrounds are mismatched. Whole *Wolbachia* genome sequencing of the wAlbB^Q^ transinfection revealed very few changes compared to the reference genome. Our results indicate that wAlbB infections may remain stable in their effects in divergent mosquito populations and across time.

## Methods

### Strain production

#### Mosquito strains and colony maintenance

*Aedes aegypti* mosquitoes were reared in temperature-controlled insectaries at 26 ± 1 °C with a 12 hr photoperiod according to Ross *et al*. (2017a). All populations were maintained at a census size of 400 individuals in BugDorm-4F2222 (13.8 L) or BugDorm-1 (27 L) cages (Megaview Science C., Ltd., Taichung, Taiwan). Larvae were reared in trays filled with 4L of reverse osmosis (RO) water and provided with fish food (Hikari tropical sinking wafers, Kyorin food, Himeji, Japan) *ad libitum* throughout their development. Embryonic microinjection experiments were performed in a general insectary, where female mosquitoes (5-7 d old, starved for 24 hr) were fed on the forearm of a human volunteer. Blood feeding of female mosquitoes on human volunteers was approved by the University of Melbourne Human Ethics Committee (approval 0723847). All adult subjects provided informed written consent (no children were involved). All following experiments were conducted in a quarantine insectary; female mosquitoes were fed human blood *via* Hemotek® membrane feeders (Hemotek Ltd, Blackburn Lancashire, Great Britain) according to Paris *et al*. (2018). Human blood was sourced from the Red Cross (Agreement #16-10VIC-02) and refreshed monthly.

Several *Wolbachia*-infected and uninfected populations were used in this study (Table 1). The *w*AlbB infection originated from *Ae. albopictus* collected in Texas, USA (Sinkins et al. 1995). *w*AlbB was introduced to *Ae. aegypti* through embryonic microinjection (Xi *et al*. 2005b) and repeatedly backcrossed to an Australian nuclear background (Axford et al. 2016). We then introduced *w*AlbB into *Ae. aegypti* with an Australian mitochondrial haplotype and nuclear background (QL) through embryonic microinjection (see below). We have named this transinfection *w*AlbB^Q^ based on its mtDNA of Queensland origin.

**Table 1.** Summary of *Aedes aegypti* populations used in this study.

| Name | Origin | *Wolbachia* infection | Mitochondrial haplotype (see Table S2) | Nuclear background (estimated %) | Replicate populations | Experiments included in |
| --- | --- | --- | --- | --- | --- | --- |
| wAlbB Au | This study | wAlbB<sup>Q</sup> | 1 | Australian - Au (100%) | 2 | All phenotypic assays, genomics |
| wAlbB My | This study | wAlbB<sup>Q</sup> | 1 | Malaysian - My (~87.5%),<br>Australian - Au (~12.5%) | 2 | All phenotypic assays |
| Uninfected Au | This study | Uninfected (tetracycline-cured) | 1 | Australian - Au (100%) | 2 | All phenotypic assays |
| Uninfected My | This study | Uninfected (tetracycline-cured) | 1 | Malaysian - My (~87.5%),<br>Australian - Au (~12.5%) | 2 | All phenotypic assays |
| wAlbB US | XI *et al.* (2005b);<br>AXFORD *et al.* (2016) | wAlbB | 3 | Australian - Au (~100%) | 1 | Microinjection only |
| QL | Collected from Cairns, Australia in 2018 | Uninfected (native) | 1, 4, 5 | Australian - Au (100%) | 1 | Microinjection and backcrossing only |
| MY | ANT *et al.* (2018) | Uninfected (native) | 2 | Malaysian - My (100%) | 1 | Backcrossing only |
| wMel | WALKER *et al.* (2011) | wMel | 1 | Australian - Au (100%) | 1 | Heat stress only |

#### Embryonic microinjection

To generate *w*AlbB-infected *Ae. aegypti* with an Australian mitochondrial haplotype (*w*AlbB^Q^), we transferred *w*AlbB from a donor population (Xi *et al*. 2005b; Axford et al. 2016) into uninfected Australian *Ae. aegypti* (QL). We used two approaches to obtain *Wolbachia* from the donor population. In the first approach, approximately 10 pairs of ovaries were dissected from females blood fed 4-5 d earlier and crushed gently with a pestle in SPG buffer in a 1.7 mL Eppendorf tube, then processed according to Xi and Dobson (2005). In the second approach, cytoplasm was removed from donor eggs and injected directly into the recipient embryo (Xi *et al*. 2005a).

For all experiments, recipient eggs were collected by placing cups filled with larval rearing water and lined with filter paper (diameter 90 mm) into cages of mosquitoes that were blood fed 4-5 days earlier. Filter papers were replaced every 0.5-1 h. Eggs (<1.5 hr old, light gray in color) were lined up on filter paper and transferred to a cover slip with double-sided tape. Eggs were left to desiccate for 1-2 min before being covered in halocarbon oil 700 (Sigma Aldrich, Castle Hill, NSW Australia). Eggs were injected using a MINJ-1000 microinjection system (Tritech Research, Los Angeles, CA, USA) and left in oil for 2 hr. Eggs were gently removed from the oil using a fine paintbrush, rinsed in water and placed on a moist piece of filter paper. Eggs were conditioned at ∼80% humidity then hatched 3 d post-injection by submerging filter papers in containers filled with RO water, a few grains of yeast and one tablet of fish food. Hatching larvae were reared to adulthood and F_0_ females were crossed to males from an uninfected population (QL), blood fed, isolated for oviposition, and screened for *w*AlbB infection after producing viable offspring. This process was repeated for three further generations, with only progeny from *Wolbachia*-positive females contributing to the next generation. The population was closed (with no further backcrossing) once the *w*AlbB infection reached fixation. We used LAMP assays for rapid detection of *w*AlbB during microinjection experiments according to Jasper *et al*. (2019).

#### Antibiotic curing

To generate uninfected *Ae. aegypti* mosquitoes with matching mitochondrial haplotypes and nuclear backgrounds, we cured the wAlbB infection using antibiotic treatment, with untreated populations reared in parallel. *w*AlbB Au (generation 5 post-microinjection) adults were fed 2 mg/mL tetracycline hydrochloride (≥95%, Sigma-Aldrich P/L, Castle Hill NSW, Australia) in a 10% sucrose solution for 10 days before blood feeding. Larvae from the next generation were reared in a solution of 50µg/L tetracycline hydrochloride according to Endersby-Harshman et al. (2019). This process was repeated for a total of three generations of adult treatment and two generations of larval treatment. After the third generation of adult treatment, 150 females from the treated population were blood fed and isolated for oviposition, with larvae from *Wolbachia-*negative mothers pooled to generate uninfected populations. To control for potential effects of drift or inbreeding (Ross *et al*. 2019a), *w*AlbB-infected (untreated) and uninfected (tetracycline-cured) females from each population were crossed to wild-type uninfected males of Australian background for one generation. The resulting populations were divided into two replicate populations for backcrossing (Figure 1A).

**Figure 1.**
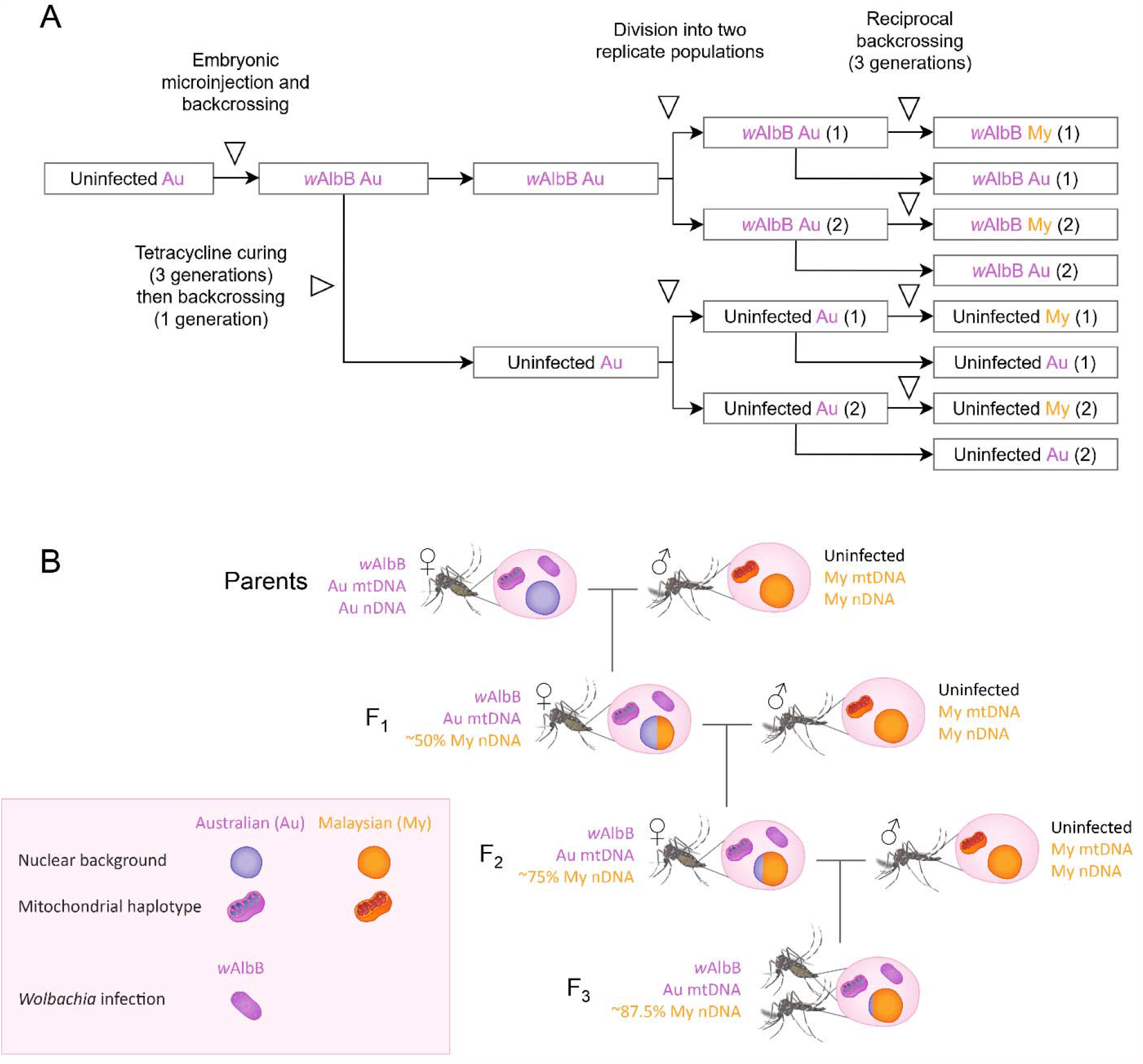
Experimental design and backcrossing scheme. (A) Experimental design showing the division of populations before and after tetracycline curing and before backcrossing. (B) Backcrossing scheme showing the introgression of the *w*AlbB infection and Australian (Au) mitochondria into a Malaysian (My) nuclear background. *w*AlbB Au females were crossed to uninfected Malaysian (MY) males for three consecutive generations. Backcrossing was also conducted with Uninfected Au females into a Malaysian background. Mosquito illustrations by Ana Ramírez (2018).

#### Backcrossing

To generate populations with different combinations of nuclear background (Australian or Malaysian) and *Wolbachia* infection status, we performed backcrosses with an uninfected Malaysian population (Figure 1B). *w*AlbB Au and Uninfected Au females were crossed to uninfected Malaysian (MY) males. Two replicate populations for each combination were backcrossed independently for a total of four populations. Each generation, 200 female pupae and 200 male pupae were sexed and left to emerge in separate cages. Sexes were confirmed following adult emergence and females were aspirated into cages of males to mate. This process was repeated for two additional generations, resulting in an estimated 87.5% similarity to the target background. Four additional populations (two replicates each of *w*AlbB Au and Uninfected Au) were not backcrossed. Instead, these were maintained at a census size of 400 individuals each generation. Before experiments commenced, all populations were screened for *w*AlbB infection as well as their COI haplotype (see below) to check for contamination during antibiotic curing and backcrossing.

### DNA level characterization of Wolbachia

#### Whole genome sequencing

We sequenced the whole *Wolbachia* genome of the *w*AlbB^Q^ transinfection generated in this study after 11 generations post-transinfection. Genomic DNA was extracted from a pooled sample of 5 individuals using a DNeasy Blood & Tissue kit (Qiagen, Hilden, Germany). Extracted DNA was randomly fragmented to a size of 350bp then end-polished, A-tailed, and ligated with Illumina sequencing adapters using a NEBNext® Ultra™ DNA Library Prep Kit (New England Biolabs, Ipswich MA, USA), and further PCR enriched with primers of P5 and P7 oligos. The PCR products which comprised the final libraries were purified (AMPure XP system, Beckman Coulter Life Sciences, Indianapolis IN, USA) and subjected to quality control tests that included size distribution by Agilent 2100 Bioanalyzer (Agilent Technologies, CA, USA), and molarity measurement using real-time PCR. The libraries were then pooled and sequenced on a NovaSeq 6000 (Illumina) using 2 × 150 bp chemistry by Novogene HK Company Limited, Hong Kong.

#### Reference genome assembly

Quality filtering of sequencing reads was performed with Trimmomatic (Bolger *et al*. 2014), with the following parameter settings: leading = 3; trailing = 3; slidingwindow = 4:15; minlen= 71. Reads were aligned to a *w*AlbB reference genome (Sinha *et al*. (2019), GenBank accession CP031221.1) using the Burrows-Wheeler Aligner (Li 2013), with the bwa mem algorithm and default parameter settings. Subsequent quality filtering and variant calling was performed with SAMtools and BCFtools (Li *et al*. 2009; Danecek *et al*. 2021). PCR duplicates were excluded from downstream analyses by soft masking. Reads with a MAPQ score < 25 were removed from the alignment, except for reads with MAPQ = 0, which were permitted to allow for mapping to repetitive regions. Genomic likelihoods were calculated using a maximum of 2000 reads per position. For variant calling, ploidy was set to haploid. The variant call output was used to create a consensus nucleotide sequence, wherein genome positions with coverage < 5 were masked as ‘N’. The consensus sequence was compared to the reference genome and two additional wAlbB genomes, originating from Hainan, China and Florida, USA (Kulkarni *et al*. 2019b; Kulkarni *et al*. 2019a). Polymorphic nucleotide positions were identified by aligning the genome sequences with Geneious v 9.1.8 (https://www.geneious.com). Regions of the Hainan and Florida *w*AlbB genomes displaying anomalous patterns of nucleotide variation, suggestive of localized sequence contamination, were annotated by eye.

#### Phylogenomic analysis

Genome assemblies and corresponding protein annotation data were obtained from GenBank (Appendix S1). Thirteen representative supergroup B genomes from various arthropod hosts were included in the analysis, including three publicly available *w*AlbB genomes (Kulkarni *et al*. 2019a; Kulkarni *et al*. 2019b; Sinha *et al*. 2019) and two supergroup A outgroup genomes (Appendix S1). Orthofinder v2.5.2 (Emms and Kelly 2019) was used to identify a core set of orthologs that were present as single-copy genes in all of the *Wolbachia* genomes included in the analysis (Appendix S2). Ortholog nucleotide sequences were aligned with MAFFT v7.475 (Katoh and Standley 2013) using the ‘auto’ setting. Alignments were trimmed with Gblocks v0.91b (Castresana 2000) using the following settings: minimum number of sequences for a conserved position = seqs/2 + 1, minimum number of sequences for a flank = seqs/2 + 1, maximum number of contiguous non-conserved positions = 8, minimum block length = 5, gap allowed = none. Sixteen ortholog alignments were excluded from subsequent analyses due to possible sequence contamination in one sample (CP041923.1; see above). For phylogenetic analysis, the individual orthogroup alignments were concatenated to form a single alignment comprised of 459 orthogroups and 382,520 nucleotide positions. Maximum likelihood trees were constructed with RAxML-HPC v8.2.12 (Stamatakis 2014) on XSEDE, using a GTR-GAMMA model with rapid bootstrapping (100 inferences). Bootstrap scores were plotted onto the best scoring ML tree. RAxML-HPC was accessed through the CIPRES Science Gateway (Miller *et al*. 2010).

#### COI sequencing

To confirm the successful transfer of the wAlbB infection to *Ae. aegypti* with an Australian mitochondrial haplotype, we performed COI sequencing on six individuals from the original *w*AlbB population described in Xi et al. (2005b) as well as nine individuals from the *w*AlbB Au population generated through microinjection in the current study. To check for contamination following backcrossing, we performed COI sequencing from three individuals from each of the 8 populations used in experiments (Table 1). We performed COI amplicon sequencing following Yeap *et al*. (2016). Samples were analysed for a 750bp region within the COI region (positions 1994-2743 on GenBank: EU352212.1) using forward primer UEA5 5’AGTTTTAGCAGGAGCAATTACTAT3’ and reverse primer UEA10 5’TCCAATGCACTAATCTGCCATATTA3’ (Lunt *et al*. 1996). PCR amplicons from individuals were sequenced in both forward and reverse directions using Sanger Sequencing (Macrogen, Inc., Geumcheongu, Seoul, South Korea). The sequenced 750bp region was analysed using Geneious 9.1.8 to investigate SNP variation among samples.

#### Wolbachia detection and density

We used real-time PCR assays (Lee et al. 2012; Axford et al. 2016) to confirm the presence or absence of *Wolbach*ia infection and estimate relative density using the Roche LightCycler® 480. Genomic DNA was extracted using 250 μL of 5% Chelex 100 Resin (Bio-Rad laboratories, Hercules CA) and 3 μL of Proteinase K (20 mg/mL) (Roche Diagnostics Australia Pty. Ltd., Castle Hill New South Wales, Australia). Tubes were incubated for 60 min at 65°C then 10 min at 90°C. Three primer sets were used to amplify markers specific to mosquitoes, *Ae. aegypti* and *w*AlbB. For mosquitoes carrying the *w*Mel infection, *Wolbachia* density was determined using w1 primers (Lee e*t al*. 2012). Relative Wolbachia densities were determined by subtracting the crossing point (Cp) value of the *Wolbachia*-specific marker from the Cp value of the mosquito-specific marker. Differences in Cp were averaged across 2-3 consistent replicate runs, then transformed by 2^n^. For the maternal transmission experiment we required only presence-absence data, so a single run was performed per sample (with any negative samples repeated to confirm a lack of infection).

### Experimental comparisons

#### Life history parameters

To evaluate the effects of *Wolbachia* infection status and nuclear background on mosquito life history, we performed phenotypic assessments of the 8 populations generated through backcrossing (Table 1). Eggs (<1 week old) from each population were hatched in trays filled with 3L of RO water, a few grains of yeast, and one tablet of fish food. One day after hatching, 100 larvae were counted into trays filled with 500 mL of RO water, with 5 replicate trays per population. Larvae were provided with fish food *ad libitum* until pupation. To determine the average larval development time and survival to pupation for each tray, pupae were counted and sexed twice per day (in the morning and evening). Pupae were removed from larval development trays, pooled across replicate trays, and placed in an open container of water inside a BugDorm-4F2222 (13.8 L) cage for adults to emerge.

Groups of adults were stored within 24 hr of emergence for *Wolbachia* density and wing length measurements. One wing each from 20 males and 20 females per population was dissected and measured for length according to Ross *et al*. (2016) as an estimate of body size, with damaged wings excluded from analysis. Adults from the four *w*AlbB-infected populations (20 females and 20 males per population) were screened with real-time PCR for *Wolbachia* density estimation (as described above). The remaining adults were provided with a 10% sucrose solution which was then removed 24 hr prior to blood feeding. Females (6-7 d old) were fed human blood via Hemotek® membrane feeders. Twenty engorged females per population were isolated for oviposition in 70 mL specimen cups with mesh lids that were filled with 15 mL of larval rearing water and lined with a strip of sandpaper (Norton Master Painters P80 sandpaper, Saint-Gobain Abrasives Pty. Ltd., Thomastown, Victoria, Australia). Eggs were collected four days after blood feeding, partially dried, and then hatched three days after collection by submerging sandpaper strips in containers filled with RO water, a few grains of yeast and one tablet of fish food. Female fecundity was determined by counting the total number of eggs on each sandpaper strip, while hatch proportions were determined by dividing the number of hatched eggs (with a clearly detached egg cap) by the total number of eggs per female. Females that did not lay eggs or died before laying eggs were excluded from the analysis.

#### Quiescent egg viability

Blood-fed females remaining in cages from the previous experiment were used to test quiescent egg viability. Six cups filled with larval rearing water and lined with sandpaper strips were placed inside each cage. Eggs were collected five days after blood feeding, partially dried, then placed in a sealed chamber with an open container of saturated potassium chloride solution to maintain a constant humidity of ∼84%. When eggs were 1, 4, 7, 10, 13 and 16 weeks old, small sections of each sandpaper strip were removed and submerged in water with a few grains of yeast to hatch. Four to six replicate batches of eggs were hatched per population at each time point, with 40-125 eggs per batch. Hatch proportions were determined by dividing the number of hatched eggs (with a clearly detached egg cap) by the total number of eggs per female.

#### Cytoplasmic incompatibility and Wolbachia density following heat stress

We measured *Wolbachia* density and cytoplasmic incompatibility in adults after being exposed to cyclical heat stress during the egg stage. Eggs were collected from *Wolbachia*-infected populations (one replicate population each from *w*AlbB Au, *w*AlbB My and *w*Mel). Four days after collection, batches of 40-60 eggs were tipped into 0.2 mL PCR tubes (12 replicate tubes per population) and exposed to cyclical temperatures of 29-39°C for 7 d in Biometra TProfessional TRIO 48 thermocyclers (Biometra, Göttingen, Germany) according to Kong *et al*. (2016) and Ross *et al*. (2019b). Eggs of the same age from each population, as well as Uninfected Au eggs, were kept at 26°C. Eggs held at 29-39°C and 26°C were hatched synchronously and larvae were reared at a controlled density (100 larvae per tray of 500 mL water). Pupae were sexed and 15 males and 15 females per population and temperature treatment were stored in absolute ethanol within 24 hr of emergence for *Wolbachia* density estimation (see *Wolbachia* detection and density). The remaining pupae were sexed and released into BugDorm-4S1515 (5.4 L) cages (with each sex, temperature treatment and population held in separate cages) for cytoplasmic incompatibility crosses.

We established two sets of crosses to 1. test the ability of *Wolbachia*-infected males to induce cytoplasmic incompatibility and 2. test the ability of *Wolbachia*-infected females to restore compatibility with *Wolbachia*-infected males. In the first set, Uninfected Au females (untreated) were crossed with *Wolbachia-*infected males from each temperature treatment by aspirating females into cages of males. In the second set, *Wolbachia*-infected females from each temperature treatment were crossed with *w*AlbB Au males, except for *w*Mel females which were crossed to *w*Mel (untreated) males. Females (5-7 d old, starved for 24 hr) were blood-fed and 20 females per cross were isolated for oviposition. Hatch proportions were determined by dividing the number of hatched eggs (with a clearly detached egg cap) by the total number of eggs per female. Females that did not lay eggs or died before laying eggs were excluded from the analysis.

#### Wolbachia maternal transmission

We tested the fidelity of *Wolbachia* maternal transmission in each nuclear background. Females from the *w*AlbB Au and *w*AlbB My populations (one replicate population each) were crossed to Uninfected Au males. Females (5-7 d old, starved for 24 hr) were blood-fed, isolated for oviposition, then stored in absolute ethanol after laying eggs. Offspring from each female were hatched, reared in trays of 500 mL RO water, then stored 4 d after hatching. We tested ten mothers and ten offspring per mother for the presence of *w*AlbB using real-time PCR (see *Wolbachia* detection and density).

#### Statistical analysis

All data were analyzed using SPSS Statistics version 24.0 for Windows (SPSS Inc, Chicago, IL). Data sets were tested for normality with Shapiro-Wilk tests and transformed where appropriate (egg hatch proportion and survival to pupa data were logit transformed). Life history, quiescent egg viability and *Wolbachia* density data were analyzed with general linear (mixed effect) models (GLMs). We tested the effects of *Wolbachia* infection (*w*AlbB-infected or uninfected), nuclear background (Au or My) and interactions between *Wolbachia* infection and nuclear background. Replicate populations were pooled for analysis when effects of replicate population exceeded a P-value of 0.1 in prior analyses. Where interaction terms were not-significant, the terms were dropped and the models were rerun without interactions. Data for each sex were analyzed separately. For quiescent egg viability, hatch proportions differed substantially between *w*AlbB-infected and uninfected populations. We therefore ran separate GLMs for *w*AlbB-infected and uninfected populations, with egg storage duration included as a continuous covariate for this trait. Replicate population (nested within nuclear background) was included as a random factor but was not significant in any instance. Quadratic egg storage duration was also included as a factor in the GLM in case of a non-linear relationship between egg hatch proportion and storage duration in these populations. We used chi-squared tests to determine whether sex-ratios for each population differed significantly from 1:1. Bonferroni corrections were performed on P values when multiple traits were measured in the same experiment.

For *Wolbachia* density, untransformed data (i.e. differences in Cp between *Wolbachia* and mosquito markers, before 2^n^ transformation) were used for analyses to test for effects of nuclear background and the temperature treatment (26 or 26-39°C) as a factor. We were unable to perform direct comparisons between *w*Mel and *w*AlbB strains due to using different markers for each strain; we therefore excluded *w*Mel from the overall analysis but presented it graphically. Egg hatch proportions from cytoplasmic incompatibility crosses were analysed with Mann-Whitney U tests.

## Results

### Genomic analysis of the wAlbB^Q^ transinfection

We sequenced the whole *Wolbachia* genome of the *w*AlbB^Q^ transinfection generated in this study. Its genome is almost identical to the *w*AlbB reference genome (Sinha *et al*. 2019), differing by only four single nucleotide variants (SNVs), suggesting that few genetic changes have occurred since *w*AlbB was first transferred to *Ae. aegypti* over 15 years ago (Xi *et al*. 2005b). The genomes of the *w*AlbB variants from Hainan, China and Florida, USA were more divergent, each differing from the *w*AlbB reference genome by more than 100 SNPs, substitutions and small indels, and by the presence of at least one large deletion (>100 bp) for the Hainan variant and at least four large deletions for the Florida variant (Appendix S3).

We performed a phylogenomic analysis with a core set of protein-encoding gene ortholog sequences conserved as single-copy genes across various *Wolbachia* supergroup B infections, including four *w*AlbB variants and two supergroup A outgroup strains. The *w*AlbB variants formed a monophyletic group that shared a most recent common ancestor with the *Wolbachia* strains *w*No, *w*Mau and *w*Tpre from Drosophila simulans, *Drosophila mauritiana and Trichogramma pretiosum* respectively (Figure 2A). A separate analysis of only the *w*AlbB sequences placed the *w*AlbB^Q^ transinfection in a cluster with the *w*AlbB reference variant, in agreement with the above sequencing results (Figure 2B). Relative to the *w*AlbB reference and *w*AlbB^Q^ transfection genomes, the Hainan variant was slightly less divergent than the Florida variant, consistent with the pattern of nucleotide variation observed for the whole genome sequences (Appendix S3).

**Figure 2.**
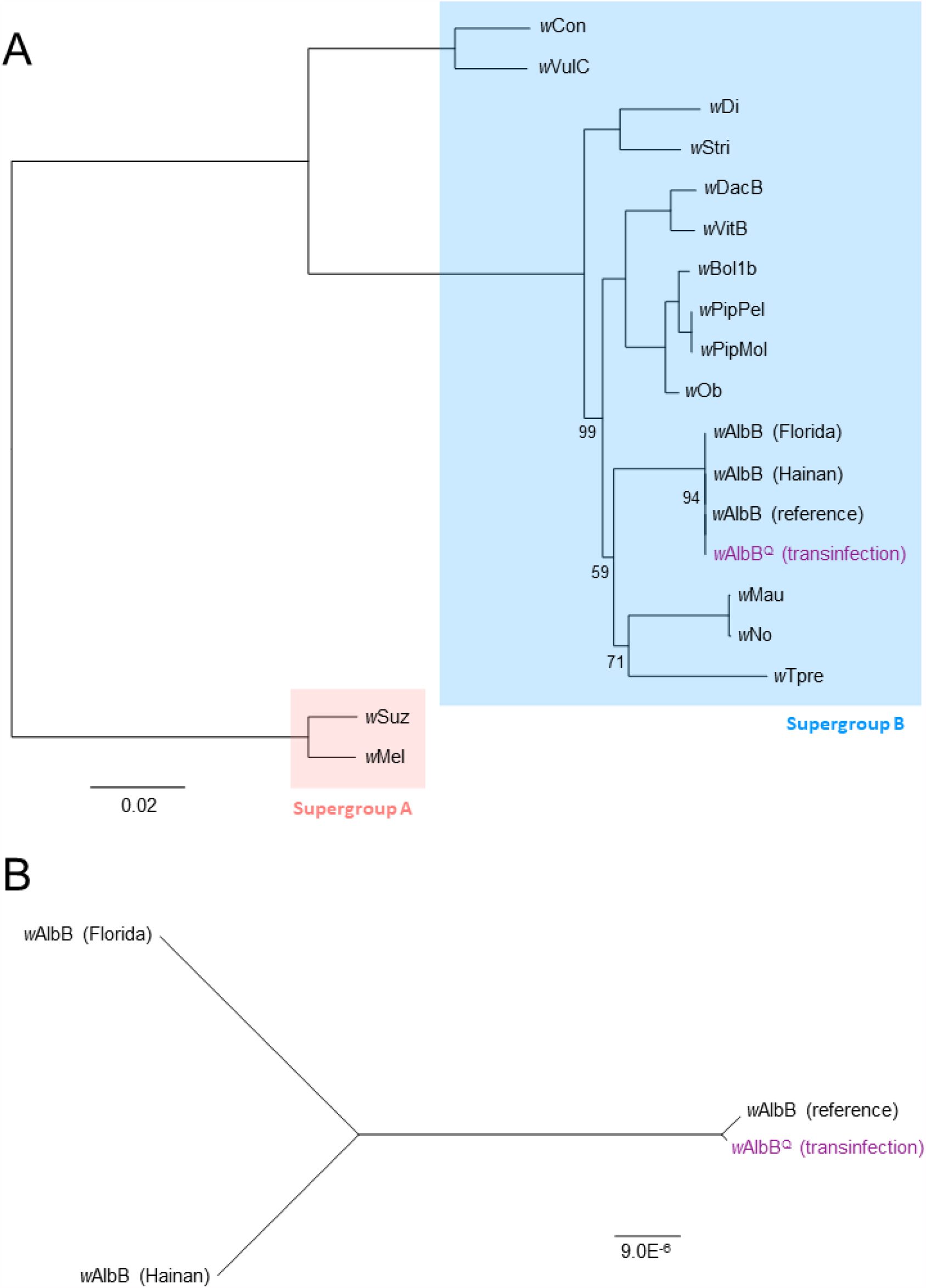
Phylogenomic analyses of *w*AlbB variants. Maximum likelihood trees were constructed with RAxML-HPC using a concatenated alignment of 459 gene orthologs, comprised of 382,520 nucleotide positions. Bootstrap values are 100 unless shown otherwise; scale bars = number of substitutions per site. The wAlbB^Q^ transinfection is highlighted in purple. For the analysis of (A) *Wolbachia* from different host species, the alignment comprised 9,469 distinct patterns and a rooted tree was built using two supergroup A sequences as outgroups; for the analysis of (B) *w*AlbB variants, an unrooted tree was built using an alignment with 23 distinct patterns.

### Life history traits are influenced by *w*AlbB infection and nuclear background

To test the contributions of *Wolbachia* infection and nuclear background to mosquito life history, we generated 8 divergent populations using a combination of embryonic microinjection, antibiotic curing and backcrossing (Table 1, Figure 1). *w*AlbB was successfully introduced to mosquitoes with an Australian mitochondrial haplotype through cytoplasm transfer, with the resulting transinfection (wAlbB^Q^) showing stable transmission by the third generation (Table S1). Following backcrossing, all 8 populations had COI sequences matching their expected mitochondrial haplotype (Table 1, Table S2).

Cohorts of mosquitoes from each population were evaluated for life history traits including larval development time, survival to pupa, fecundity, egg hatch proportion and body size (Figure 3). Development time and survival to pupa were not significantly affected by *Wolbachia* infection or nuclear background (Table 2). Nuclear background had a substantial effect on fecundity (Table 2), with Malaysian background mosquitoes laying more eggs than those with an Australian background (Figure 3E). The *w*AlbB infection reduced the proportion of eggs hatching, but there was no significant effect of nuclear background (Table 2, Figure 3F). Mosquitoes with a Malaysian background tended to have larger wings than mosquitoes with an Australian background (Figure 3G, H), but this effect was only significant in males (Table 2). Sex ratios of pupae (Figure 3D) did not deviate significantly from 1:1 (Chi-square: df = 9, all P > 0.05) for any population. Overall, our results indicate clear effects of nuclear background on fecundity and wing length, with *w*AlbB infection inducing a cost to egg hatch in both nuclear backgrounds.

**Table 2.** GLMs for life history parameters of *w*AlbB-infected and uninfected *Aedes aegypti* of Australian (Au) and Malaysian (My) backgrounds. P values in bold are significant following Bonferroni correction (Adjusted α: 0.00556).

| Trait | Factor | Type III Sum of Squares | Degrees of freedom | Mean Square | F | P |
| --- | --- | --- | --- | --- | --- | --- |
| Female development time | *Wolbachia* infection | 0.016 | 1 | 0.016 | 1.279 | 0.265 |
|  | Nuclear background | 0.085 | 1 | 0.085 | 6.729 | 0.014 |
|  | Error | 0.468 | 37 | 0.013 |  |  |
| Male development | *Wolbachia* infection | 0.032 | 1 | 0.032 | 1.175 | 0.285 |
| time |  |  |  |  |  |  |
|  | Nuclear background | 0.123 | 1 | 0.123 | 4.571 | 0.039 |
|  | Error | 0.995 | 37 | 0.027 |  |  |
| Survival to pupa | *Wolbachia* infection | 0.003 | 1 | 0.003 | 1.379 | 0.248 |
|  | Nuclear background | 0.000 | 1 | 0.000 | 0.177 | 0.677 |
|  | Error | 0.075 | 37 | 0.002 |  |  |
| Fecundity | *Wolbachia* infection | 5.663 | 1 | 5.663 | 0.020 | 0.889 |
|  | Nuclear background | 4573.419 | 1 | 4573.419 | 15.874 | **< 0.0001** |
|  | Error | 42351.708 | 147 | 288.107 |  |  |
| Egg hatch proportion | *Wolbachia* infection | 0.595 | 1 | 0.595 | 10.911 | **0.001** |
|  | Nuclear background | 0.060 | 1 | 0.060 | 1.096 | 0.297 |
|  | Error | 7.856 | 144 | 0.055 |  |  |
| Female wing length | *Wolbachia* infection | 0.007 | 1 | 0.007 | 0.920 | 0.339 |
|  | Nuclear background | 0.034 | 1 | 0.034 | 4.538 | 0.035 |
|  | Error | 1.163 | 157 | 0.007 |  |  |
| Male wing length | *Wolbachia* infection | 0.001 | 1 | 0.001 | 0.377 | 0.540 |
|  | Nuclear background | 0.088 | 1 | 0.088 | 27.554 | **< 0.0001** |
|  | Error | 0.544 | 171 | 0.003 |  |  |

**Figure 3.**
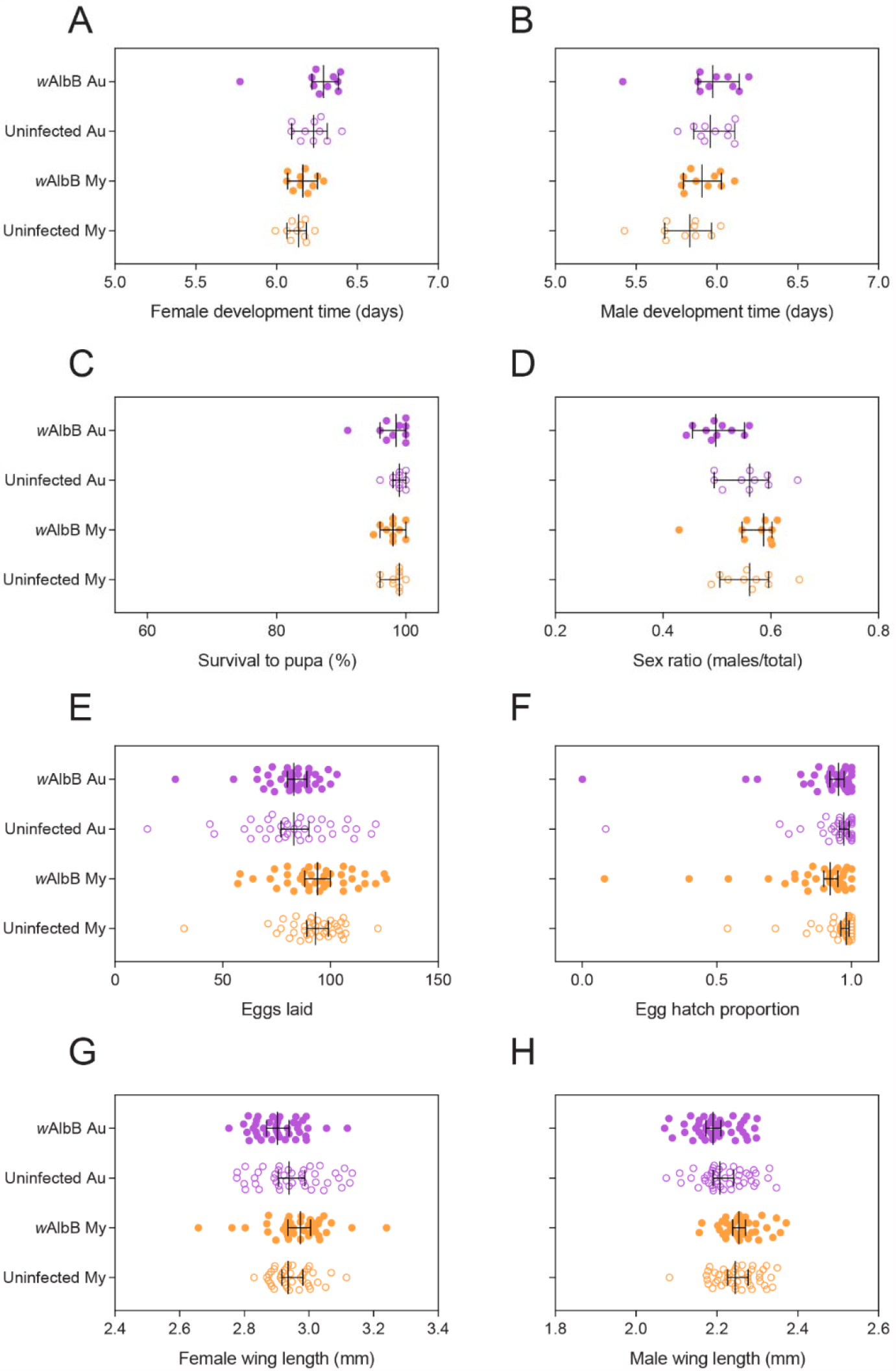
Life history parameters of *w*AlbB-infected and uninfected *Aedes aegypti* of Australian (Au) and Malaysian (My) backgrounds. Populations were evaluated for the following traits: larval development time for (A) females and (B) males, (C) percent survival to the pupal stage, (D) sex ratio, (E) female fecundity, (F) egg hatch proportion and wing length of (G) female and (H) males. Data from two replicate populations were pooled for visualization. See table 1 for a description of each population. Each point represents data averaged across a replicate container of 100 individuals (A-D) or data from individual mosquitoes (panels E-H). Medians and 95% confidence intervals are shown in black lines.

### Quiescent egg viability depends on mosquito nuclear background and *w*AlbB infection

Stored eggs from each population were hatched every three weeks to determine quiescent egg viability. wAlbB infection greatly reduced quiescent egg viability in both Australian and Malaysian backgrounds (Figure 4). By week 16, hatch proportions for *w*AlbB-infected populations approached zero while hatch proportions for uninfected populations exceeded 40%. In uninfected populations, a LM indicated a significant interaction between background and week (F_(1, 118)_ = 13.193, P < 0.001) due to the sharp decrease in viability in the Malaysian background. There was also an effect of week in this analysis (F_(1, 118)_ = 98.967, P < 0.001). Eggs with an Australian background had higher hatch proportions (median 0.819) than eggs with a Malaysian background (median 0.445) by the end of the experiment. In wAlbB-infected populations where we did not consider the data from week 16, there was also a significant interaction between background and week (F_(1, 93)_ = 22.948, P < 0.001) due to the sharper decrease in viability in the Malaysian background along with an effect of week (F_(1, 93)_ = 368.012, P < 0.001).

**Figure 4.**
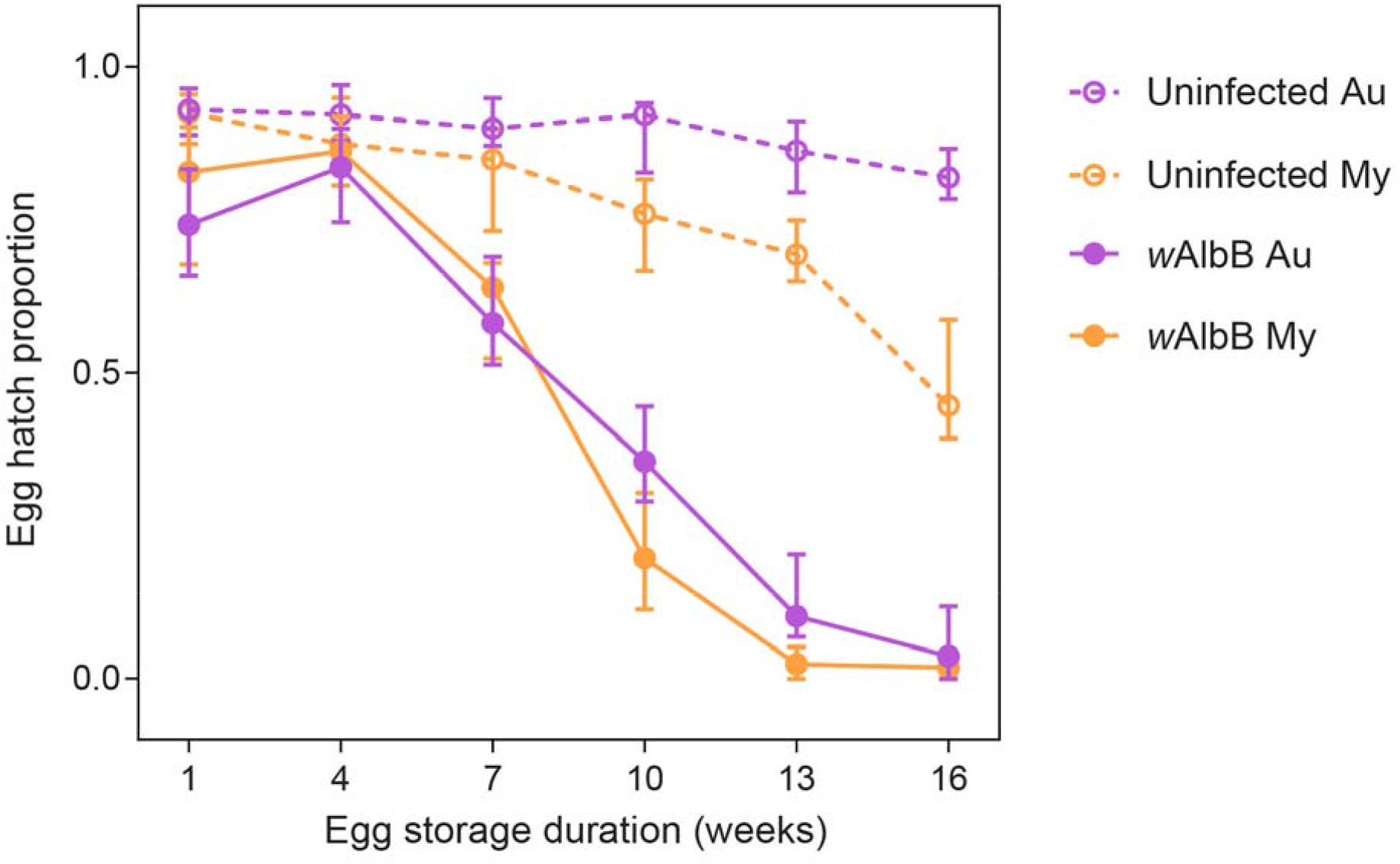
Quiescent egg viability of *w*AlbB-infected and uninfected *Aedes aegypti* of Australian (Au) and Malaysian (My) backgrounds. Data from two replicate populations were pooled for visualization. Symbols show median egg hatch proportions while error bars show 95% confidence intervals.

### Nuclear background influences *w*AlbB density

We estimated *Wolbachia* density in whole adults from the *w*AlbB Au and *w*AlbB My populations (Figure 5). *Wolbachia* density was higher in Australian mosquitoes than Malaysian mosquitoes for both females (GLM: F_1,82_ = 6.752, P = 0.011) and males (F_1,72_ = 7.956, P = 0.006).

**Figure 5.**
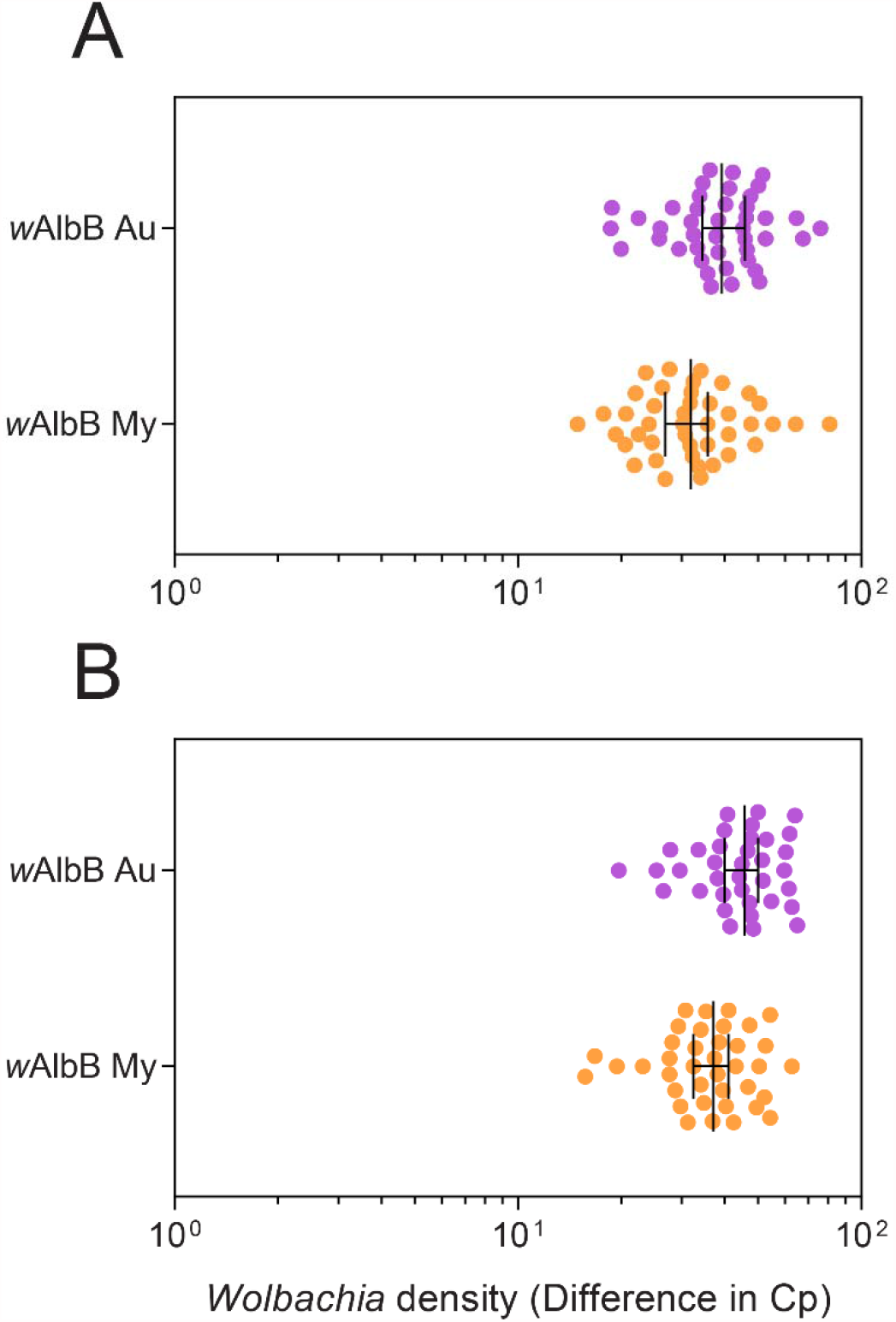
Female (A) and male (B) *Wolbachia* density in *w*AlbB-infected *Aedes aegypti* of Australian (Au) and Malaysian (My) backgrounds. Data from two replicate populations were pooled for visualization. Each point represents the relative density for an individual averaged across 2-3 technical replicates. Medians and 95% confidence intervals are shown in black lines.

### *w*AlbB shows complete maternal transmission regardless of nuclear background

We observed complete maternal transmission of wAlbB in both the Australian and Malaysian nuclear backgrounds, with all 10 offspring from 10 females (100/100) in each population being infected (lower 95% confidence interval: 0.96).

### *w*AlbB is stable under cyclical heat stress regardless of nuclear background

To test the stability of *w*AlbB at high temperatures in the different nuclear backgrounds, we measured *Wolbachia* densities in adults after eggs were exposed to cyclical heat stress (29-39°C) or held at 26°C. We found no significant effect of temperature (females: F_(1, 56)_ = 2.073, P = 0.155, males: F_(1, 56)_ = 0.676, P = 0.414), nuclear background (females: F_(1, 56)_ = 2.228, P = 0.141, males (F_(1, 56)_ = 1.341, P = 0.252), or interactions between temperature and nuclear background (females: F_(1, 56)_ = 1.276, P = 0.264, males: F_(1,56)_ = 0.749, P = 0.391), indicating that this infection is stable under heat stress regardless of nuclear background (Figure 6). In contrast, the *w*Mel infection decreased in density by a median of 98.7% (across both sexes) under the same conditions (Figure 6).

**Figure 6.**
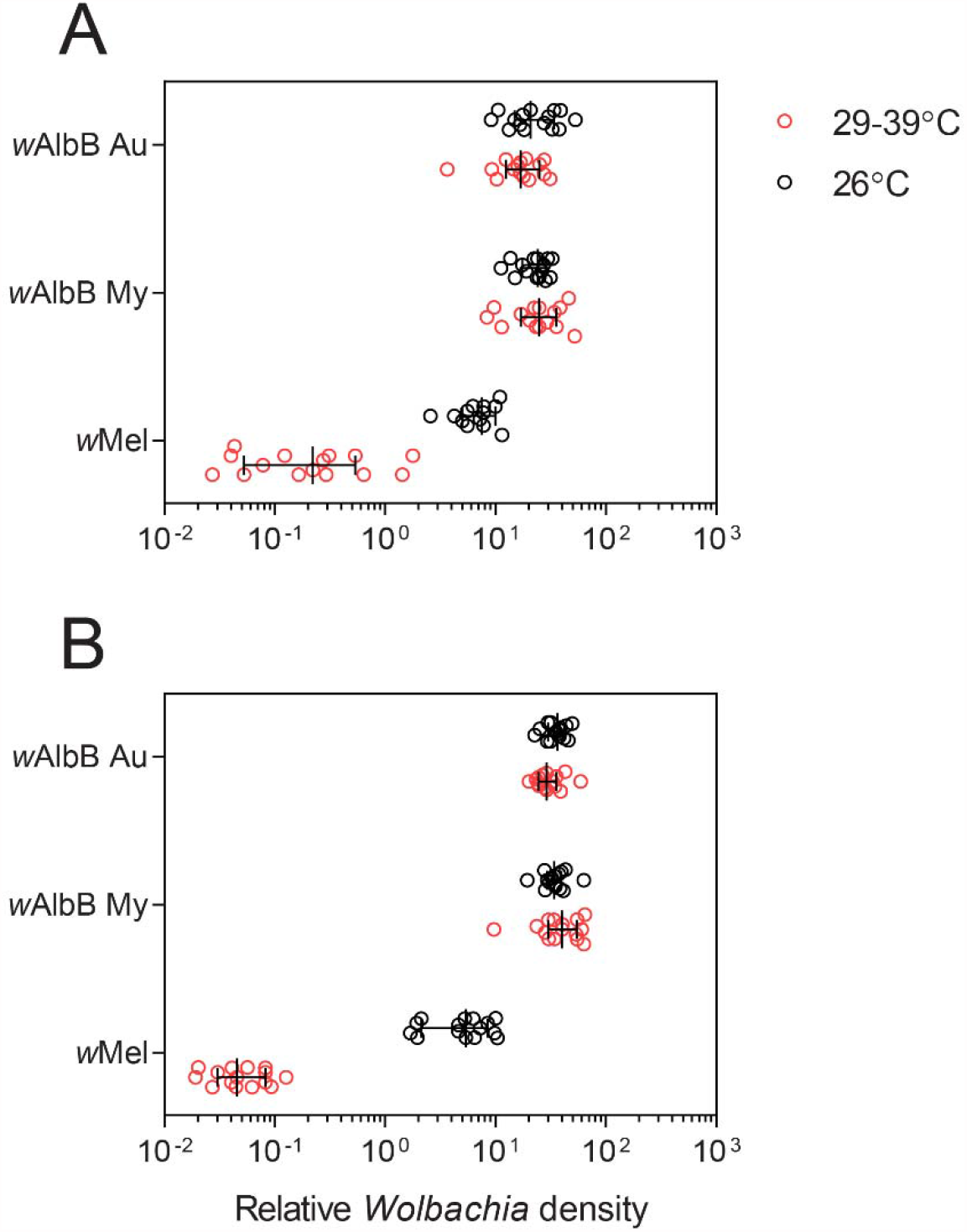
*Wolbachia* density in (A) females and (B) males following exposure to cyclical heat stress during the egg stage. Eggs from the *w*AlbB Au, *w*AlbB My and *w*Mel populations were exposed to cyclical temperatures of 29-39°C for 7 d (red circles) or held at 26°C (black circles). Each point represents the relative density for an individual averaged across 2-3 technical replicates. Medians and 95% confidence intervals are shown in black lines.

### *w*AlbB induces complete cytoplasmic incompatibility regardless of nuclear background and heat stress

We tested the ability of *w*AlbB-infected males to induce cytoplasmic incompatibility. Uninfected females that were crossed to *w*AlbB Au or *w*AlbB My males produced no viable offspring (no eggs hatched), indicating complete cytoplasmic incompatibility (Figure 7A). *w*AlbB-infected males continued to induce complete cytoplasmic incompatibility following exposure to cyclical temperatures of 29-39°C for 7 d during the egg stage, regardless of nuclear background. In contrast, heat stress weakened cytoplasmic incompatibility induction by *w*Mel-infected males, with all uninfected females in this cross producing viable offspring (Figure 7A).

**Figure 7.**
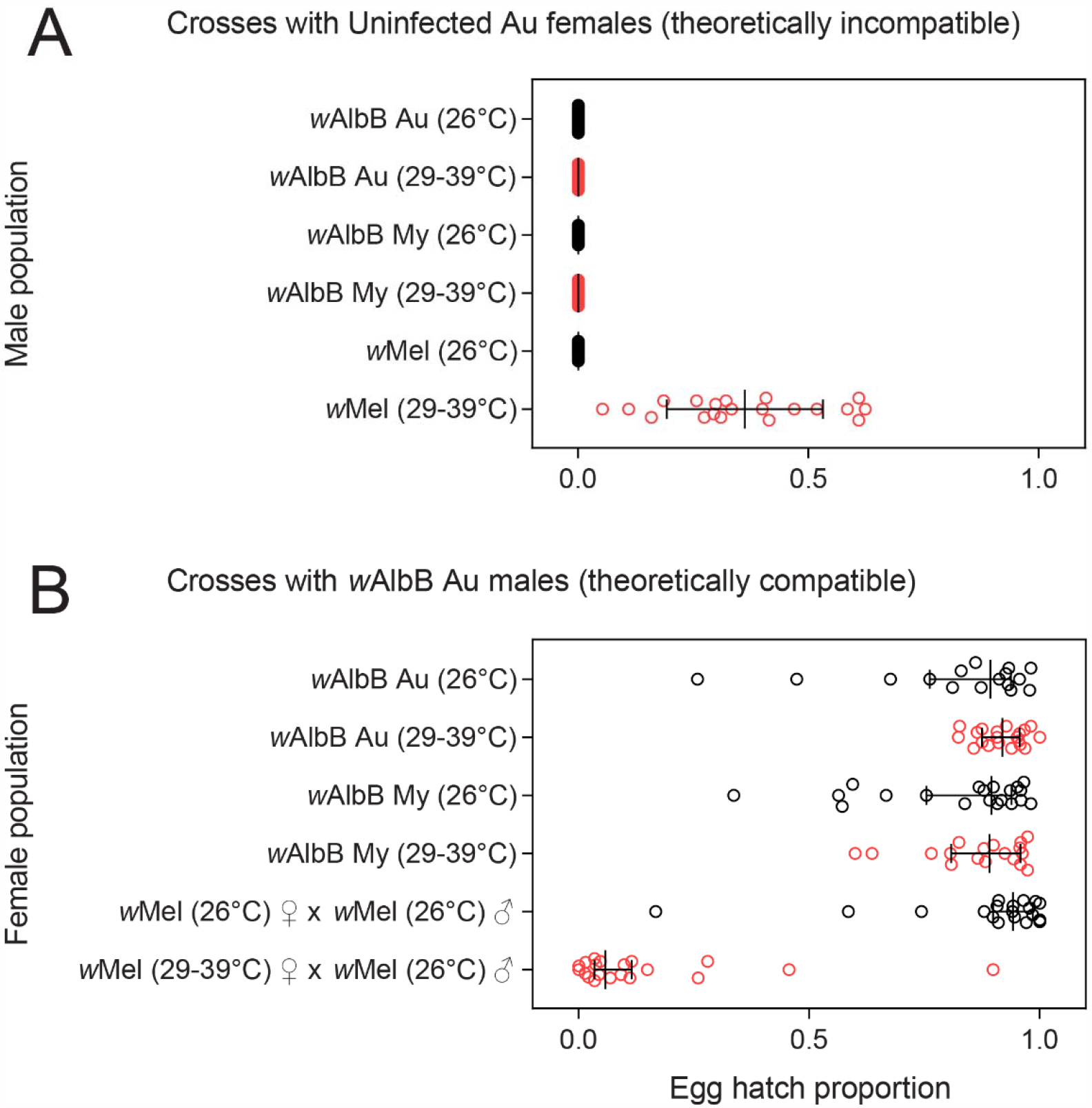
Cytoplasmic incompatibility induction and compatibility restoration following exposure to cyclical heat stress during the egg stage. We performed crosses to test the ability of (A) *Wolbachia-* infected males to induce cytoplasmic incompatibility with uninfected females and (B) *Wolbachia*-infected females to restore compatibility with *Wolbachia*-infected males. In each cross, males (A) or females (B) from the *w*AlbB Au, *w*AlbB My and *w*Mel populations were exposed to cyclical temperatures of 29-39°C for 7 d (red circles) or held at 26°C (black circles) during the egg stage. Each point represents the hatch proportion of eggs from a single female. Medians and 95% confidence intervals are shown in black lines.

We also tested the ability of *w*AlbB-infected females to restore compatibility with *w*AlbB-infected males under heat stress (Figure 7B). *w*AlbB-infected females exhibited high hatch proportions regardless of nuclear background or heat stress. Heat stress had no significant effect on the egg hatch proportions of *w*AlbB Au (Mann-Whitney U: Z = 1.512, P = 0.131) or *w*AlbB My females (Z = 0.585, P = 0.562), but greatly reduced the egg hatch of *w*Mel-infected females (Z = 5.114, P < 0.001). Overall, these results show that both cytoplasmic incompatibility induction and compatibility restoration are robust to nuclear background and heat treatment.

## Discussion

In this study, we generated a wAlbB transinfection (*w*AlbB^Q^) in *Ae. aegypti* and evaluated its phenotypic effects in two nuclear backgrounds. *w*AlbB showed 100% maternal transmission, induced complete cytoplasmic incompatibility and was stable at high temperatures across both Australian and Malaysian backgrounds. Life history traits were driven largely by mosquito nuclear background, with Malaysian mosquitoes having larger wings and laying more eggs than Australian mosquitoes, but with lower quiescent egg viability. However, *w*AlbB infection induced substantial costs to egg hatch, particularly in quiescent eggs. These effects were consistent across two replicate populations. We found no clear deleterious effects of having mismatched mitochondrial and nuclear genomes, supporting the use of backcrossing as a method to introduce *Wolbachia* infections into target backgrounds for field release. Whole *Wolbachia* genome sequencing of the *w*AlbB^Q^ transinfection revealed very few changes compared to the reference genome. Our results point to the stability of *w*AlbB infections across time, environment, and host background. The *w*AlbB^Q^ transinfection generated in this study is therefore likely to retain desirable traits for arbovirus and mosquito control under a broad range of conditions.

*w*AlbB infections in *Ae. aegypti* typically induce host fitness costs (Axford *et al*. 2016; Joubert *et al*. 2016; Ant et al. 2018; Lau *et al*. 2021), but these effects may depend on both host and *Wolbachia* variation. Carvalho *et al*. (2020) recently demonstrated the importance of host background, where *w*AlbB showed larger deleterious effects in a Mexican background compared to a Brazilian background. Here we found that *w*AlbB infection reduced egg hatch proportions and dramatically reduced quiescent egg viability, but effects were consistent across two mosquito backgrounds. Although the backgrounds used in Carvalho et al. (2020) were different from ours, differences in outcomes could also be explained by methodological differences. We used antibiotic curing to ensure that mitochondrial haplotypes were matched between infected and uninfected lines, while Carvalho et al. (2020) compared the *w*AlbB-infected populations to naturally-uninfected populations. However, we also used fewer rounds of backcrossing, resulting in incomplete introgression into the Malaysian background. Controlling for mitochondrial haplotype is an important consideration when elucidating *Wolbachia* phenotypes given that it can have substantial effects on host fitness in other insects (Hoekstra *et al*. 2013; Meiklejohn *et al*. 2013; Rank *et al*. 2020).

We performed whole genome sequencing of our *w*AlbB^Q^ transinfection to detect potential evolutionary changes and found that its sequence is almost identical to the reference genome (Sinha *et al*. 2019). The transinfection and reference genomes have a common origin; both *w*AlbB infections were derived from *Ae. albopictus* caught in Houston, Texas, USA in 1986 (Sinkins et al. 1995). However, their histories are distinct; the reference sequence was derived from a wAlbB infection maintained in the Ae. aegypti Aa23 cell line for more than two decades (O’Neill *et al*. 1997), while the transinfection was introduced to *Ae. aegypti* over 15 years ago (Xi et al. 2005b). The long-term stability of wAlbB is consistent with the *w*Mel and *w*MelPop-CLA infections in *Ae. aegypti*, which also show few long-term genomic (Woolfit et al. 2013; Huang et al. 2020) and phenotypic (Hoffmann et al. 2014; Ross *et al*. 2020a) changes following transinfection. These results suggest that *Wolbachia* infections will likely remain stable in their effects following field releases, broadly consistent with phenotypic data for a separate *w*AlbB transinfection following releases in Malaysia (Ahmad *et al*. 2021).

The *w*AlbB *Wolbachia* strain is widespread throughout natural populations of *Ae. albopictus* but is thought to exhibit low genetic diversity (Bourtzis *et al*. 2014). Previous surveys have detected little or no genetic variation within *w*AlbB through a limited set of molecular markers (Armbruster *et al*. 2003; Albuquerque *et al*. 2011; Das et al. 2014; Hu *et al*. 2020), but a recent reanalysis of *Ae. albopictus* genome sequence data points to substantial variation within this strain (Scholz et al. 2020). We compared the genome sequence of our *w*AlbB^Q^ transinfection to two other publicly available sequences from *Ae. albopictus* caught in St. Augustine, Florida, USA in 2016 (Kulkarni et al. 2019a) and Haikou, Hainan, China in 2016 (Kulkarni et al. 2019b) and found that all three were distinct from each other. These results emphasize the need to reevaluate *w*AlbB diversity in *Ae. albopictus*, since variation within wAlbB could influence control interventions targeting this species. This also raises the question of whether independently-derived *w*AlbB transinfections show phenotypic differences. For instance, the *w*AlbB transinfection generated by Xi *et al*. (2005b) is stable at high temperatures (Ross *et al*. 2017b) while the transinfection generated by Ant *et al*. (2018) that has been deployed in Malaysia (Nazni *et al*. 2019) decreases in density under a similar temperature regime (Ant et al. 2018). Given the sequence variation within *w*AlbB, nomenclature for differentiating between variants (as for other *Wolbachia* strains such as *w*Pip (Atyame et al. 2011)) may be useful, particularly if they differ in their phenotypic effects.

*Aedes albopictus* is native to Asia and only invaded USA in the 1980s (Benedict *et al*. 2007). The initial incursion into mainland USA is thought to have originated in Japan, and to have been followed by a complex pattern of successive introductions from other locations (Zhong *et al*. 2013; Goubert *et al*. 2016; Kotsakiozi *et al*. 2017; Manni *et al*. 2017). Given that the Houston strain of *Ae. albopictus*, from which the *w*AlbB^Q^ transinfection and the reference variant are derived, was collected only one year after the first report of *Ae. albopictus* in mainland USA in 1985 (Sprenger and Wuithiranyagool 1986), it is likely that this strain originated in Japan. Recent population genetic studies of *Ae. albopictus* have identified clear patterns of geographic differentiation among populations within the native Asian range of the species (Kotsakiozi *et al*. 2017; Schmidt *et al*. 2020). Consistent with this, our phylogenomic analysis separated the *w*AlbB variants into three main branches: a Chinese branch, a Florida Branch (of unknown origin) and a probable Japanese branch. Understanding the extent of variation within *w*AlbB in natural populations could provide insights into the speciation and global spread of *Ae. albopictus*.

In conclusion, we have shown that phenotypic effects of the *w*AlbB infection tested here (and associated mtDNA) are stable across nuclear backgrounds, providing little evidence for nuclear genome-*Wolbachia* interactions or changes in *w*AlbB associated with multiple host transfers through microinjection. Our results have implications for releases of *Wolbachia-*infected *Ae. aegypti* that are taking place in different mosquito backgrounds around the world. They suggest that one source of infection may serve releases in multiple locations, although adequate backcrossing remains important to ensure that the nuclear background is consistent with that of target populations for genes under local selection such as those involved with pesticide resistance (Endersby-Harshman *et al*. 2020).

## Acknowledgements

We thank Meng-Jia Lau, Véronique Paris, Moshe Jasper, Ashley Callahan, Jason Axford and Vanessa White for technical assistance and Ian Forster and Trent Perry for needle pulling assistance. We thank Zhiyong Xi for providing the original wAlbB transinfection used in this study.

## Data availability

The whole genome sequence of the wAlbB^Q^ transinfection will be deposited in Genbank following submission.

## Supplementary information

**Table S1.**
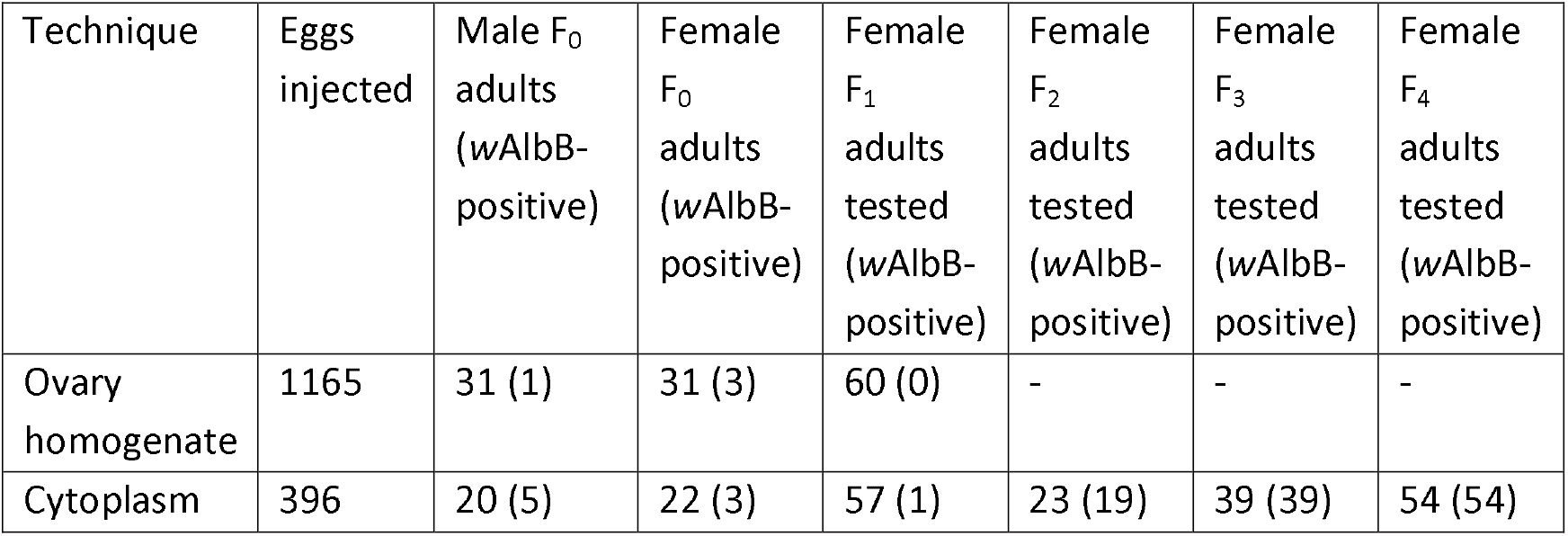
Generation of a stable *w*AlbB transinfection (*w*AlbB^Q^) in an Australian mitochondrial and nuclear background of *Aedes aegypti* through embryonic microinjection.

**Table S2.** Haplotype combinations for the mitochondrial COI region.

| Mitochondrial haplotype | SNP position (750 bp fragment) |  |  |  |  |  |  |  |  |  |  |  |
| --- | --- | --- | --- | --- | --- | --- | --- | --- | --- | --- | --- | --- |
|  | 3 | 84 | 192 | 414 | 426 | 531 | 579 | 594 | 630 | 636 | 682 | 744 |
| 1 | C | T | C | T | T | A | C | A | C | T | T | T |
| 2 | T | C | C | C | C | G | T | G | T | C | C | T |
| 3 | C | T | C | T | T | G | C | A | C | T | T | T |
| 4 | T | T | T | C | C | G | T | G | T | T | C | T |
| 5 | C | T | C | T | T | G | C | A | C | T | T | G |

**Appendix S1. Datasets used for phylogenomic analyses**.

**Appendix S2. Protein accessions for each orthogroup**.

**Appendix S3. Genomic analysis of *w*AlbB variant polymorphisms**.

